# Error Correction of SARS-CoV-2 Genomic Sequences Using a Phylogenetic Prior

**DOI:** 10.64898/2026.10.07.757202

**Authors:** Monica Arniella, Lenore Pipes, Rasmus Nielsen

## Abstract

Genotype calling of sequencing data has improved over recent years thanks to technological and statistical breakthroughs. One such advance is the use of imputation-based genotyping methods in human genetics. For haploid and mostly non-recombining organisms, a natural equivalent is to leverage phylogenetic signal for imputation-based genotype calling. SARS-CoV-2 is one such case, and the use of improved genotyping methods in this species is particularly pertinent given the well-documented systematic problems with read alignment and assembly in rapidly evolving regions of its genome, motivating the need for statistically principled methods for genotype correction that complement existing community efforts to curate sequencing data. We present a statistical framework to combine phylogenetic prior probabilities with the information from aligned reads, and demonstrate its utility in correcting consensus calls made on short-read, viral sequencing data. We derive phylogenetically-informed posterior probabilities based on genome placement in a reference phylogeny, and read likelihoods calculated from machine-reported base quality scores. We further modify the prior at each site by estimating a maximum-likelihood scaling parameter jointly with reads at all other sites. We find that the phylogenetically-informed base calling method stays highly accurate across simulated depth and error conditions. In real datasets, the method vastly improves call rate while still maintaining similar accuracy as reference-based assembly methods. In problematic genomic regions, the phylogenetically-informed method confers both accuracy and call rate gains. Accuracy and call rate do not stratify with sequencing center and primer scheme used, suggesting broad applicability across sequencing technologies and potentially diverging sequences. Our results in real and simulated data demonstrate that phylogenetically-informed genotyping correction results in more complete sequences, achieving a higher call rate without compromising accuracy. In cases of low mean depth, sparse coverage, or difficult-to-sequence regions that may still retain a low call rate, the method improves the accuracy of genotype calls.

## 1. Introduction

Despite the availability of a variety of genomic sequencing technologies, and the promise of recent advances in long-read, single molecule (third-generation) sequencing, a majority of publicly available genomic data around the world has been generated with short-read (second-generation) technologies [Hotaling et al., 2021, Chen et al., 2022]. Illumina short-read instruments, in particular, are well-established and were quickly and successfully adopted to investigate SARS-CoV-2 upon its emergence in 2019 as a novel disease-causing pathogen [Chiara et al., 2021]. However, despite the reliability and popularity of these instruments, random and systematic errors can still arise during the template amplification and sequencing-by-synthesis steps: empirical assessments of overall error rates vary from ∼0.3-1%, and in some rare instances beyond (from 1-7%), with variability both between and within institutions, instruments, protocols, and even individual experimenters [Stoler and Nekrutenko, 2021, Mitchell et al., 2020]. In the case of viral sequencing, distinguishing true variants from sequencing errors is an important problem, made even more salient by rapid evolution, resulting issues such as amplicon dropout, and the real-time demands of pandemic-scale data analysis [Arana et al., 2022, Ulhuq et al., 2023, Markov et al., 2023].

To minimize errors, genotype calling approaches typically use heuristics such as quality score and allele frequency cutoffs, artifact removal and quality filtering of reads [Kubik et al., 2021]. In SARS-CoV-2 data, problematic sites have been additionally cataloged by community curation [De Maio et al., 2020]. Additionally, genotype calling in diploids has benefited from probabilistic approaches using patterns of linkage disequilibrium (LD) to resolve calls or impute missing data [Nielsen et al., 2011]. In the case of viruses, largely non-recombinant species such as SARS-CoV-2 have an analogous route to probabilistic genotype calling, which relies on a strong phylogenetic signal instead of LD patterns. Phylogenetic probabilities have been successfully used to impute missing nucleotides in low-coverage clinical, assembled sequences [Pipes et al., 2022], and this use case has further potential to be extended as a variant calling tool.

Computational methods to correct base calling in a sequencing-technology aware manner in large scale SARS-CoV-2 data [Hunt et al., 2026] have also resulted in progress in quantifying their effect on downstream evolutionary inference [De Maio et al., 2026]. However, these efforts rely on discerning reads affected by sequencing issues, and masking positions inferred to be problematic based on the areas targeted by unreliable reads. While more systematic than curated lists of sites, these approaches still result in several uncalled stretches of the genome. Furthermore, sequencing read correction studies often focus on moderate (10x or more) sequencing depth [Kallenborn et al., 2021, Dlugosz and Deorowicz, 2024], rather than on low and ultra-low sequencing data, where the resulting lowering of call rate due to sequencing issues is bound to be most severe. Here, we implement a framework for phylogenetically-informed variant calling, evaluating its effect on accuracy and call rate, and focusing on cases of low sequencing depth. To achieve higher quality genotyping, we calculate likelihoods using all mapped reads, integrate them with phylogenetic priors calculated from the expansive global phylogeny, and compare base calls from posterior probabilities of each nucleotide, versus calls made by state-of-the-art, single reference-based assembly methods.

## 2 Approach

Given a set of sequencing reads aligned at a particular position, we can calculate the likelihood for those reads using their reported error rate. In particular, Illumina quality scores (S) encode the probability of sequencing error (E) through the relationship *S* = −10*log*(*E*). Given the error rate *e*_*i*_ derived from the quality score for read *i*, then the contribution from read *i* to the likelihood of nucleotide *a* ∈ {*A, C, T, G*} is *q*_*ai*_ = (1 − *e*_*i*_) if nucleotide *a* is the called nucleotide in read *i* (i.e. the nucleotide is correctly called), and *q*_*ai*_ = *e*_*i*_*/*3 if the nucleotide is incorrectly called.

Phylogenetically-informed base calling relies on tronko [Pipes and Nielsen, 2024], a method which can efficiently assign reads to nodes in a phylogeny using an approximate full likelihood approach. To do this, tronko calculates posterior probabilities of each nucleotide at each node of a provided reference tree, and assigns sequences to the node with the best score. For base calling purposes, one entire assembled genome serves as the sequence being placed, and the posterior probabilities of each nucleotide at the node chosen for placement are subsequently used as prior probabilities for the base calling analysis. Let the probabilities inferred from tronko be the base calling prior probabilities *p*_*a*_, *a* ∈ {*A, C, T, G*} for a particular site. Then, the posterior probability of nucleotide *a*, combining all *n* reads for this sequence position, is:

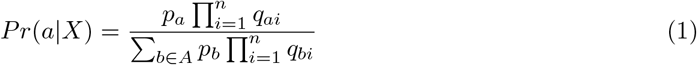

We compute posterior probabilities for each of the four nucleotides, and “call” the base as the nucleotide with the highest posterior probability. In the case where a threshold is applied, sites where no bases exceed the posterior probability threshold are called as N.

We additionally modify *p*_*a*_ to model mutational distance from the closest lineage in the reference database. First, we assume a constant scaling factor *c* for all sites, and model the effect of evolutionary changes, and other unmodeled noise, using an adjusted prior *p*_*a*_(*c*) = (*p*_*a*_ + *c/*4)*/*(1 + *c*). Let the *k*_*i*_ bases in site *i* have read likelihoods *l*_*i*1_, *l*_*i*2_, …, 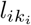 where *l*_*ij*_ = (*l*_*ijA*_, *l*_*ijC*_, *l*_*ijG*_, *l*_*ijT*_) and *l*_*ijb*_ is the read likelihood of base *b, b* ∈ *B*, in read *j* of site *i. l*_*ijb*_ can be calculated from the Phred-scaled quality scores. Then, the likelihood function for *c*, calculated for all *S* sites, is given by

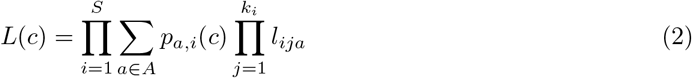

where *p*_*a,i*_(*c*) is the value of *p*_*a*_(*c*) for the *i*th site. To obtain a maximum likelihood estimate of *c, ĉ*, we optimize this function using a Golden Section Search subject to the constraint *c* ≥ 0 and we then replace *p*_*a*_ for the *i*th site with *p*_*a,i*_(*ĉ*) in Equation (1) when calculating posterior probabilities for site *i*. We can think of this as an Empirical Bayes approach where the parameter of the prior for each site is estimated using maximum likelihood jointly for all sites.

## 3 Materials and methods

For this study, we used publicly available datasets deposited in the Sequencing Read Archive (SRA) and the Global Initiative for Sharing All Influenza Data (GISAID). We also relied on the global SARS-CoV-2 multiple sequence alignment (MSA) published via GISAID on August 14, 2023.

### 3.1 Simulated sequencing reads

We downloaded fully assembled genomes from GISAID [GISAID, 2026], then simulated reads under typical error rates for Illumina instruments as follows: we queried the database for complete, high coverage, Illumina sequences, and randomly sampled genomes evenly representing the sequencing instrument models that were annotated (*n* = 69). To simulate reads, we used a custom Perl script that randomly generates reads based on a user-specified error rate per site, read length, number of reads, and insert length. We generated the per-site error rates in R by sampling from Beta distributions with shape parameters of (10, 9990), and (700, 99300), and offset the values by 1e-04 to achieve distribution means of 0.1% and 0.7% and upper and lower limits corresponding to typical error rates. We additionally generated reads under an intermediate error condition (shape parameters (40, 9960) and the same offset, mean error rate 0.4%), analyzed only in the supplementary figures.

### 3.2 Real sequencing reads

We used the query ‘(ARTIC) AND “Severe acute respiratory syndrome coronavirus 2”[Organism] AND AMPLICON[Strategy] AND VIRAL RNA[Source] AND PCR[Selection] AND PAIRED [Layout] AND Illumina[Platform] AND filetype fastq[Properties] AND (”2023/08/14”[PDAT] : “2024/08/14”[PDAT])’ to download SRA accessions deposited after the publication of the GISAID global phylogeny, selecting those for which the ARTIC primer scheme was used. We then used the SRA RunSelector to exclude non-human or cell culture samples, and further selected samples based on the “spots with mates”, “bases”, and “avgLength” fields of associated metadata to exclude sets of raw reads with low average sequencing depth. Of these, our final selection (*n* = 81) included samples with assemblies containing no N’s (after our own manual assembly), and further made sure to include samples with a complete genome deposited in GISAID. The final selection consists of sequencing runs originating from three different centers, one using an earlier primer scheme (ARTIC v2) [Merkt et al., 2023] and two using later primer schemes (ARTIC v5.3.2) [Connecticut State Public Health Laboratory, 2024, US Air Force School of Aerospace Medicine, 2023–2024].

### 3.3 Assembly and subsampling

We assembled all sets of sequencing reads with the reference-based assembly pipeline from the Broad Institute’s viral NGS suite [Broad Institute], and used the Wuhan-Hu-1 reference genome for SARS-CoV-2. For each of the full sets of raw reads, we used the pipeline’s default alignment software, minimap2 [Li, 2018], with recommended settings of skip mark dupes=true and min coverage=20, and the corresponding .bed file to trim the corresponding ARTIC adapter sequences. We set aside these high-quality, full-depth assembled genomes as the “ground-truth” for each SRA run. We then calculated the number of reads that should be sampled for each run’s set of subsamples, based on each intended mean depth and the pipeline reports of mean read length and number of bases from the ground-truth assembly. We randomly selected reads using seqtk sample [Li, 2012]. We repeated this procedure to achieve mean depths ranging from ≤ 1*x* to 20x, resulting in a set of eleven subsamples per SRA run. Each set of subsampled reads was then assembled using the same pipeline, but using min coverage=3. For the real data, we used representative subsamples to analyze base calling, retaining two per category for low (0.8x, 1x), intermediate (3x, 5x), and high (10x, 20x) sequencing depth. For the simulated sets of raw reads, we specified read length and number at the simulation step, limited the analysis to mean depths of 5x and lower, and assembled these subsamples with the same pipeline but chose to align with the BWA algorithm [Li and Durbin, 2009] due to minimap2 algorithm constraints with the simulated reads.

### 3.4 Placement in global phylogeny using tronko

We used the tronko [Pipes and Nielsen, 2024] workflow to estimate phylogenetic prior probabilities for all assembled genomes (full and subsampled depths). Specifically, tronko calculates posterior probabilities of each nucleotide at each position of the sequence being taxonomically assigned to a node in the tree; these posterior probabilities at the assigned node are then used as priors in our analyses (i.e. *p*_*a*_ as given by Equations (1) and (2), Approach). We used default parameters with tronko-assign, and placed samples with respect to a reference database using the August 14th, 2023 global MSA published through GISAID. We tested modifying tronko probabilities using different scaling factors: 0.0 for no scaling, a constant 0.001 as a proof of concept, and a varying maximum likelihood estimate (MLE) as previously described. We restricted analysis to MLE-corrected priors; the method to correct these is implemented in a custom C script.

### 3.5 Calculating posterior probabilities

We used the .bam file from the assembly pipeline outputs to generate pileup files using samtools [Li et al., 2009]. Using a custom Python script, we parsed match strings and quality scores from the pileup file to calculate the likelihoods described in equation (2), we read in the tronko probabilities described in 3.4, calculated posterior probabilities for every position that appeared in the relevant pileup, and called the nucleotide with the highest posterior probability at each position. In order to specifically assess the effect of phylogenetic error correction, we only considered positions possessing both (a) read information (i.e. nonzero reads aligned to the position in the Wuhan-Hu-1 reference) and (b) phylogenetic information (i.e. present in the global MSA, therefore excluding insertions and deletions). We did not impute missing positions, so that the accuracy of our corrected base calls can be directly compared to that of a typical reference-based read assembly. We adjusted the threshold for base calling using the posterior probability, adjusting the acceptable difference *δ* between a probability of 1 (most certain) and the actual posterior probability. We constructed the thresholds such that *δ* spans from 10^−1^ to 10^−15^, with two steps per power of 10, i.e. 1 × 10^−1^ and 5 × 10^−1^ for the power of -1, 1 × 10^−2^ and 5 × 10^−2^ for the next power, et cetera. Any bases not passing a given threshold are called as N.

### 3.6 Comparing results from different methods

We evaluated accuracy and call rate for calls made using the phylogenetically-informed posterior probability (PP) method, and calls made by a state-of-the-art (SA) reference-based assembly pipeline. To increase the chance of comparing as many of the same sites as possible across samples and sequencing centers, we also excluded any samples that had N’s in the ground-truth genome.

We define the call rate as 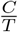, where *C* is the number of calls and *T* is the total number of sites considered (i.e. out of a total 29,903bp if genome-wide, or out of 3,831bp for the S gene). Accuracy, in turn, is defined as 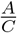, where *A* is the number of sites where the evaluated method call matched the ground-truth call, i.e. A is the total accurate sites. For accuracy, we assigned fractional values in positions where the assembly pipeline called an ambiguous base (i.e. a call of R compared to the ground-truth of A, would result in an accuracy of 0.5). These calculations were made using custom Python scripts.

We also conducted analyses either (a) over the entire genome, or (b) after restricting sites to biologically and technically relevant categories, such as sites with known poor sequencing quality [De Maio et al., 2020, UCSC Genome Browser], or sites within structural and non-structural protein-coding regions.

## 4 Results

### 4.1 Phylogenetic priors and MLE scaling improve genotyping in real and simulated reads

We evaluate the accuracy and call rate of the genotype callers using low and medium sequencing depth, mimicking data in which genotyping is difficult, as in environmental samples and other approaches that have the potential to result in low sequencing depth. For the phylogenetically-informed method (PP) method, we adjust the call rate by varying the posterior probability threshold for genotype calling.

Under both low (0.1%) and high (0.7%) simulated sequencing error rate, PP achieves higher accuracy than state-of-the-art (SA) genotyping (Fig. 1; PP (pr scaling=MLE))), when using the MLE scaling modification for the tronko priors as previously described (Methods 3.4). The unscaled prior (Fig. 1; PP (pr scaling=0.0)), that assumes no evolutionary divergence from the reference placement in the phylogeny, performs markedly worse, and has lower accuracy, for the same call rate, than both PP with scaling and the SA genotyper. In all cases there is an upper limit to the call rate as we do not call sites with 0 reads aligned, and thus no coverage. We note that the phylogenetically-informed prior has the potential to call such sites, but we have not pursued that here.

**Figure 1:**
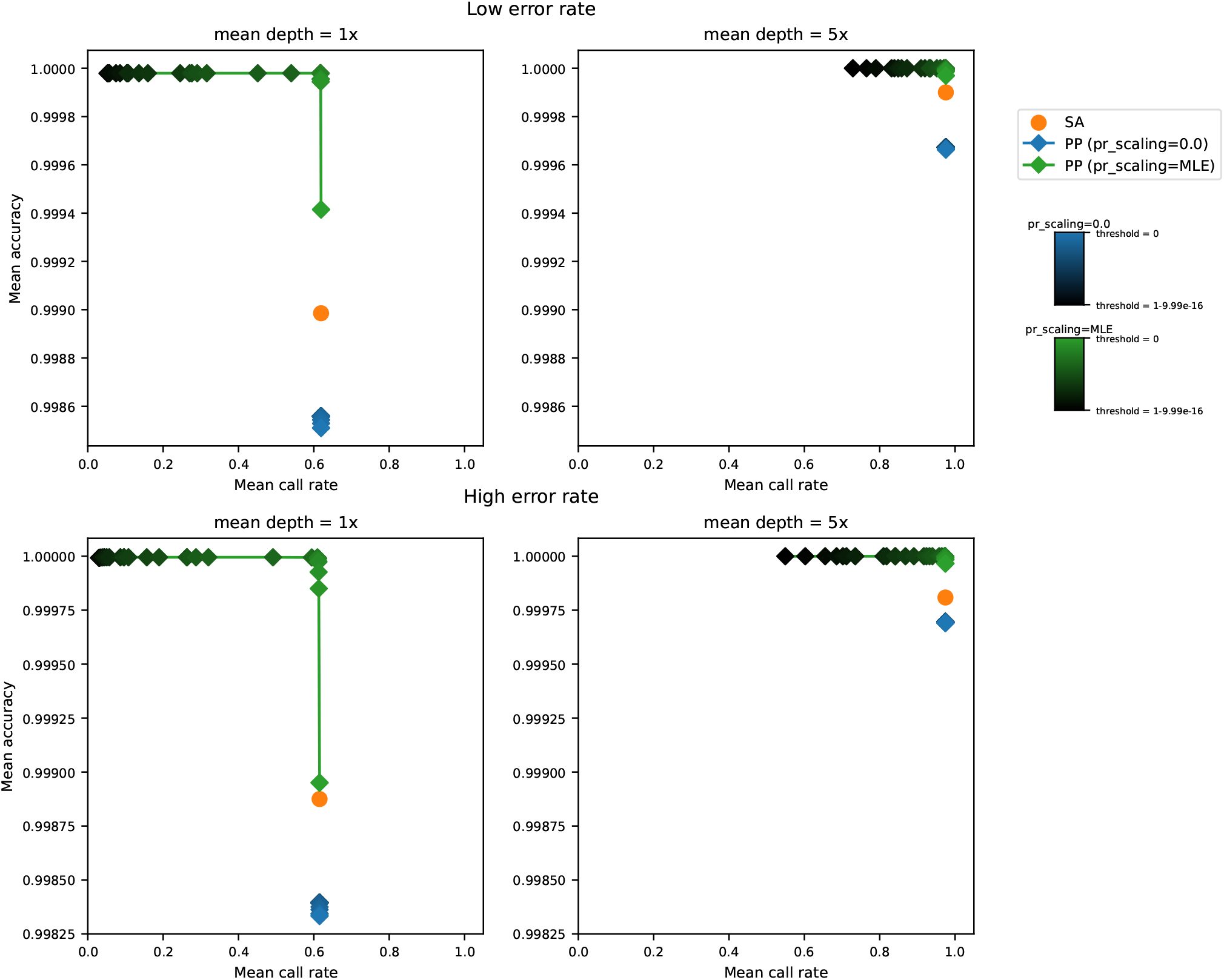
Accuracy and call rate in simulated data, at two mean genome-wide error rates. Accuracy is calculated genome-wide as the fraction of positions where the called base at a subsample (1x on the left, or 5x on the right) matches the called base at the ground truth assembly. The lowest threshold is 0, allowing all PP calls to be made regardless of certainty. Darker points denote more conservative thresholds for base-calling, resulting in an overall curve. SA calls are made using only one set of parameters, resulting in a single point. SA and PP call rate are overall high, while accuracy is higher for calls made by PP even at low thresholds.

**Figure 2:**
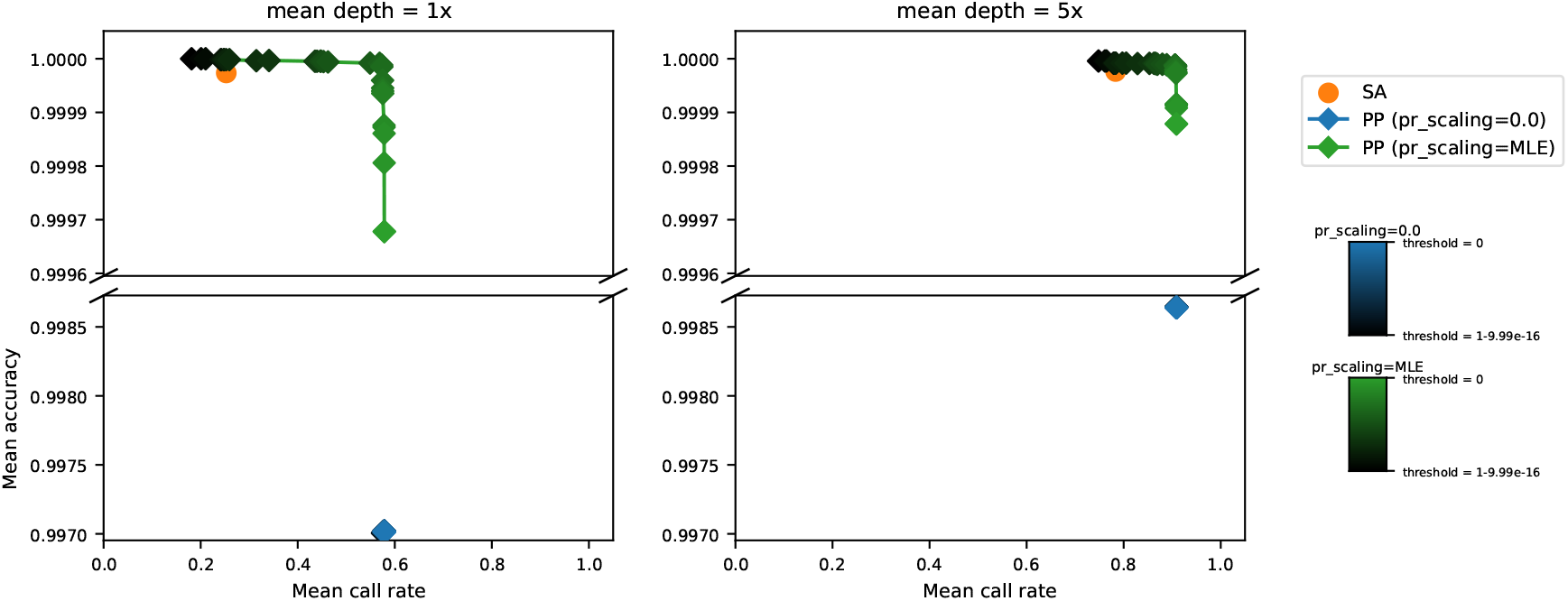
Accuracy and call rate in real data. Accuracy is calculated genome-wide as done for simulated data. We use the same PP thresholds as in Fig. 1, and interrupted axes are due to a larger gap between the accuracies of unscaled and MLE-scaled posterior probability calls. Accuracy is high for both SA and PP calls, while call rate is higher for calls made by PP, at almost all thresholds.

In real subsampled data, the SA averages reside within the convex hull of the PP curves: for example, at mean depth 1x, for the same mean call rate as SA, PP and SA have accuracies of 0.999998 and 0.999973, respectively. Similarly, calibrating PP with scaling to have the same accuracy as SA, PP and SA have call rates of 0.5733 and 0.2527, respectively (Supp. Tables S4 and S5). Thus, while applying even higher thresholds is possible, even an intermediate value will confer higher confidence base calls and dramatically increase call rate. Based on our results, a recommended threshold would be at least 1 − 10^−5^ to benefit from this improved tradeoff between accuracy and call rate (Fig. 5). As expected, the accuracy increases with sequencing depth for all methods, and the differences between methods are most apparent for subsamples at 2x and below, where traditional base-calling may not be able to resolve disagreements in the base supported by the aligned reads at an ambiguous position (Supp. Fig. S1). At 5x sequencing depth, SA is more affected by high sequencing error rates than PP with scaling, resulting in a larger difference in accuracy between SA and the least accurate PP calls made at a low threshold. It is exactly when errors cause two different nucleotides to be observed in different reads in the same position that the phylogenetic prior is important, because it helps inform which of the two observed bases is likely the correct one.

### 4.2 Site-and gene-specific challenges in genotyping SARS-CoV-2 reads

When we stratify results by quality flags from community-maintained lists of problematic sites [De Maio et al., 2020, UCSC Genome Browser], we observe a similar trend where SA falls within the PP curve regardless of site category (Fig. 3). However, “caution” flagged sites show the greatest difference at low depth, with both lower accuracy and lower call rates, though an increase in average read depth brings the two methods to similar accuracy, on par with the rest of the genome (”pass” sites). More problematic sites with the “mask” flag remain overall harder to call for both methods at any depth, with an overall lower call rate due to fewer reads aligned (Supp. Fig. S3).

**Figure 3:**
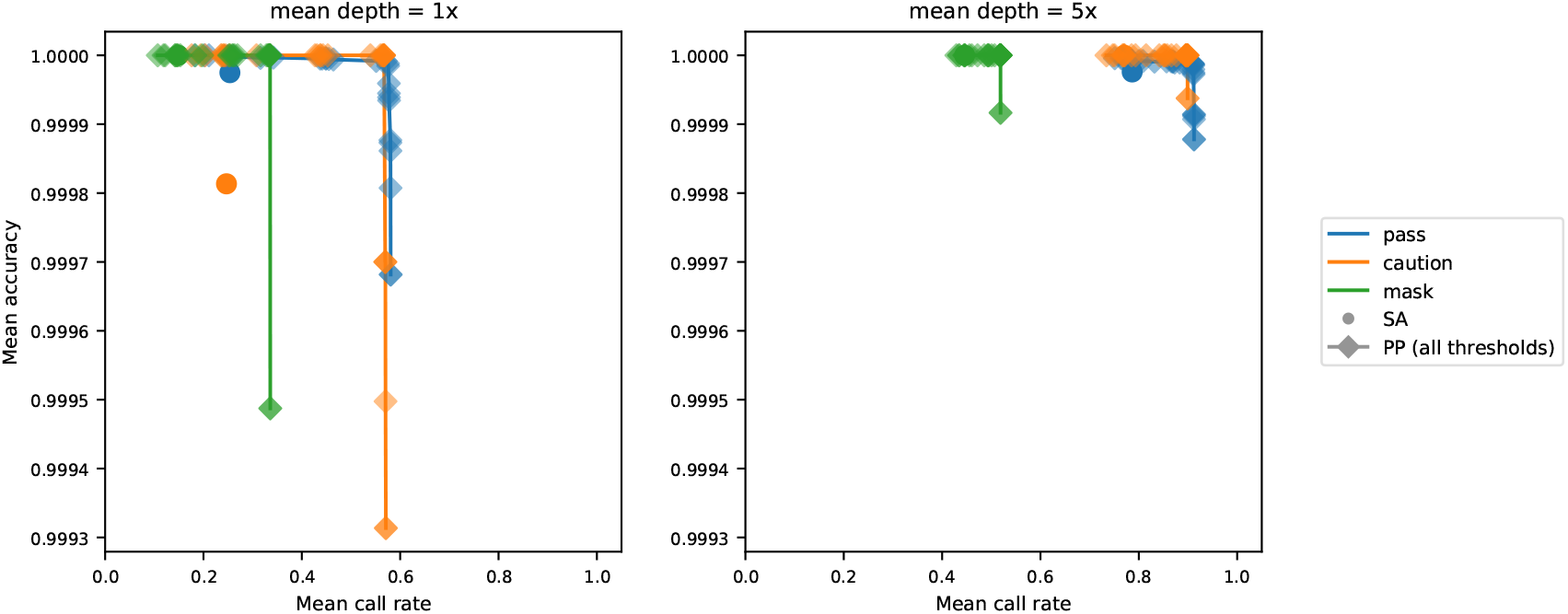
Accuracy and call rate in real data, calculated within quality flag categories of sites. The same posterior probability thresholds are used as in Figs. 1 and 2, though not shaded for clarity. Overall, PP increases call rate, though mainly at the lower mean depth, and more markedly for the “pass” and “caution” categories. All PP calls are made using MLE scaling for the prior.

Stratifying results by structural and non-structural protein regions, we see a similar trend when focusing on the S gene, which encodes the Spike protein (Fig. 4). Despite accuracy gains using the phylogenetic probability method, the overall call rate for the S gene is lower compared to other regions. This likely reflects known rapid evolution and sequencing difficulties, mirroring the circumstances of the “mask” and “caution” site categories. In contrast, the calls made at other structural and non-structural protein-coding genes have overall higher accuracy and call rates and benefit more from increased sequencing depth than do the calls made in the S gene region.

**Figure 4:**
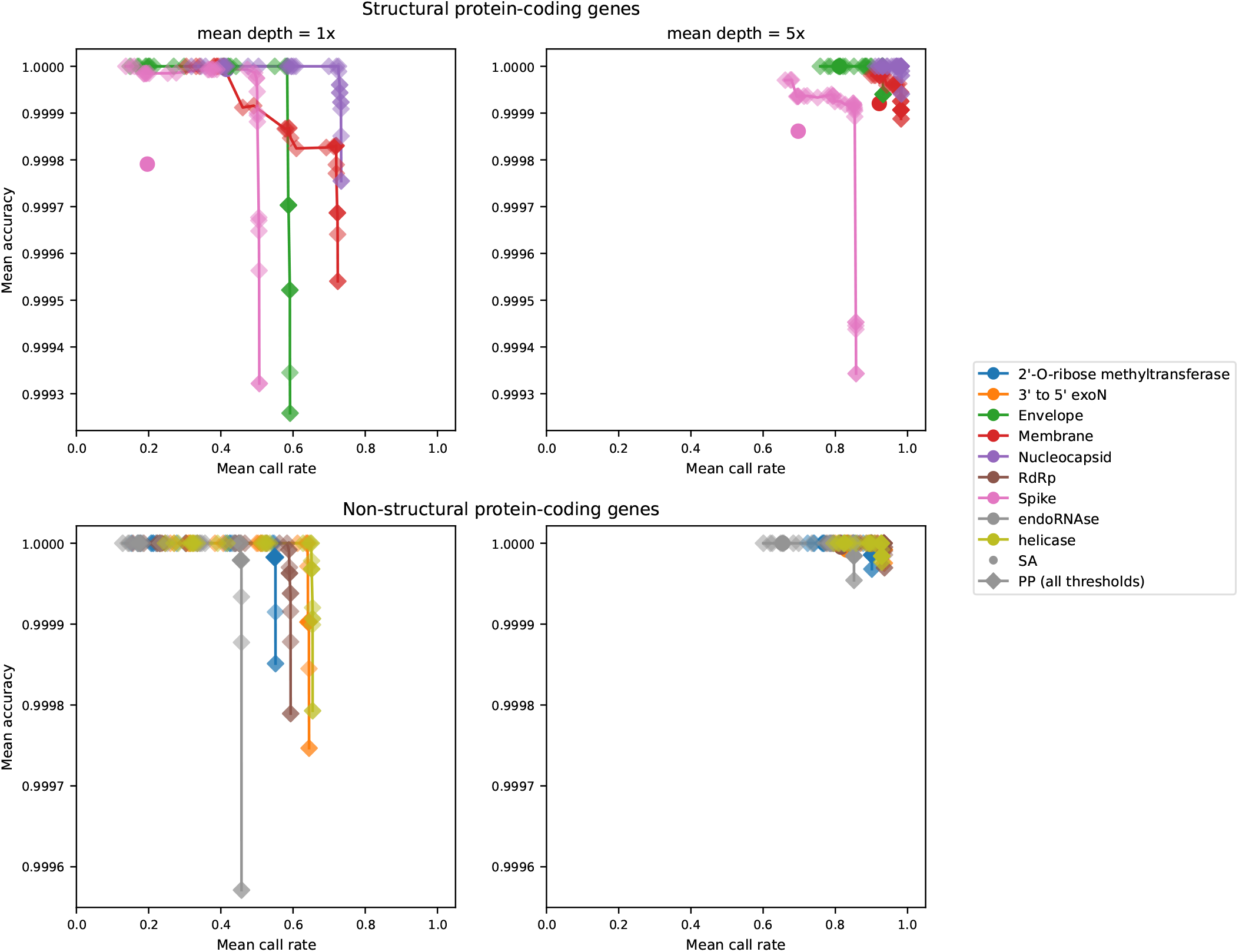
Accuracy and call rate in real data, calculated over gene regions encoding structural (top) and non-structural proteins (bottom). Overall, PP drastically increases call rate while maintaining comparable accuracy, especially at lower mean sequencing depths and at known problematic regions such as the S gene. Spike remains a region that is difficult to call for both methods, with overall lower call rates. However, PP surpasses SA in both accuracy and call rate by a wider margin. All PP calls are made using MLE scaling for the prior.

### 4.3 Center-and primer-specific analysis

When samples are grouped by sequencing center and primer, we observe the same overall gain from applying the phylogenetic probability method, with SA means within the PP curve, regardless of sequencing conditions. In aggregate and at a medium sequencing depth, we see very subtle primer-or center-specific differences that differ by depth. At mean depth of 5x, PP results are nearly indistinguishable from each other whether grouped by sequencing center or primer scheme used (Fig. 5). However, there are subtle differences in accuracy at a mean depth of 1x that fall along center differences. Specifically, results from centers that sequenced samples using a later primer scheme (ARTIC v5.3.2) show a slightly lower call rate and accuracy than those from centers that used an earlier primer scheme (ARTIC v2) (Fig. 5).

**Figure 5:**
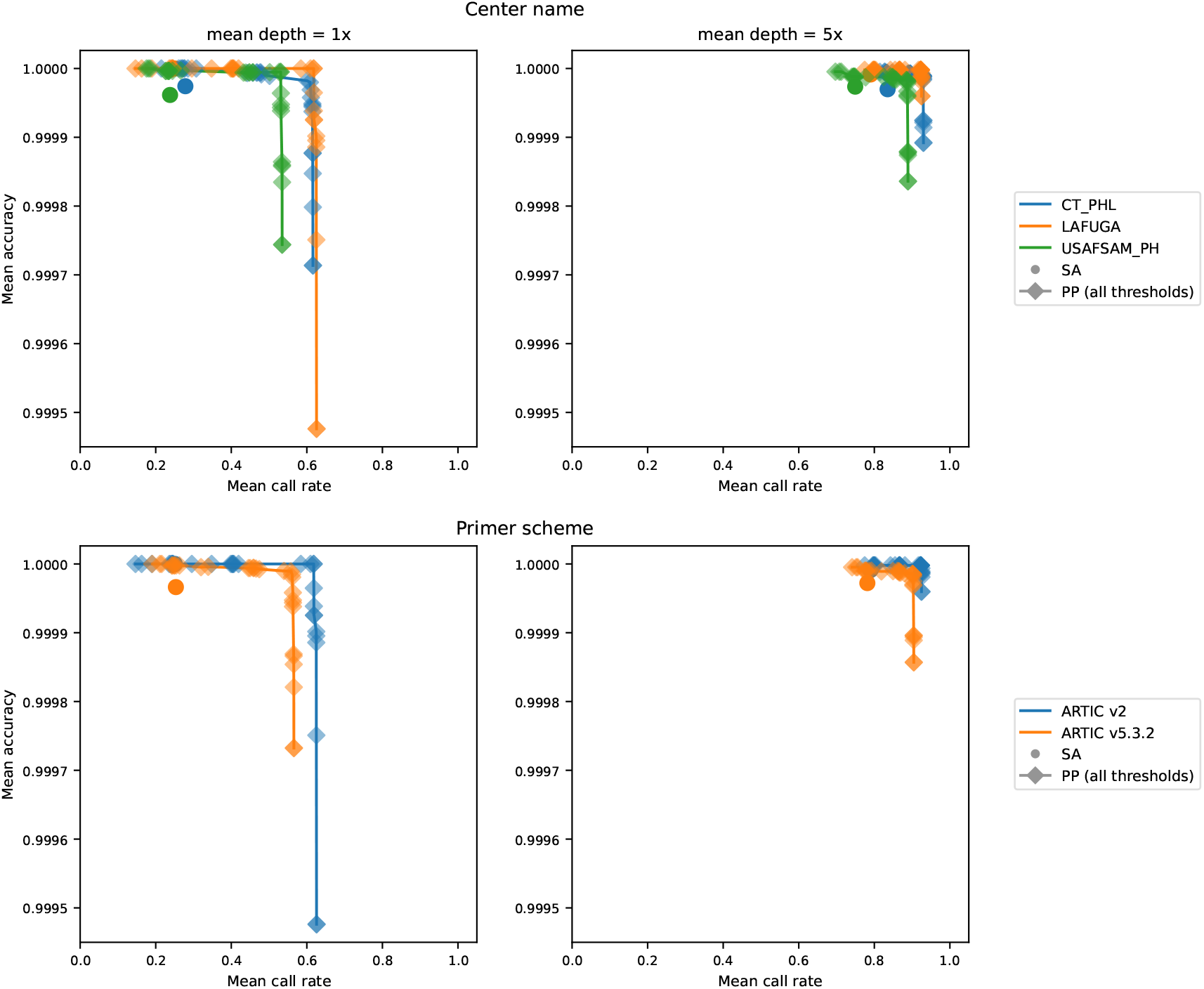
Accuracy and call rate in real data, compared between sequencing centers and primer schemes used to generate samples. All PP calls are made using MLE scaling for the prior.

When it comes to difference by sequencing centers, we may not be able to untangle sequencing batch effects (i.e. instrument, individual, or center differences) from strain-specific sequencing challenges, due to the overlap of sequencing centers with clusters of locally circulating genotypes that may be genuinely harder to sequence. Variation between individual samples is, in fact, higher than between sequencing centers (Supp. Fig. S2). This variation may partly reflect differences in the accuracy of the phylogenetic prior arising from uneven representation of sampled lineages in the reference database. Since the sequences that were generated using ARTIC v2 primers were also sampled at a timepoint before reference database construction, these samples in particular have a more accurate prior. Furthermore, at a mean depth of 1x, sparse read support may limit the ability to distinguish sequencing errors from true mutations that differ from the phylogenetic prior, particularly in lineages poorly represented in the reference database. However, errors can also arise from a mismatch between primer scheme and the strain being sequenced [Hunt et al., 2026], which can apply to earlier and later primer schemes. Even with all else being equal (in terms of the efficacy of the prior and the MLE scaling factor), sequencing errors due to primer mismatch could affect rapidly-evolving later variants the most, still differentially affecting the accuracy of the later primer scheme data. These comparisons therefore cannot isolate the effects of primer scheme from those of sampling date, lineage composition, and representation in the reference database.

## 5. Discussion

Our results demonstrate that phylogenetic prior probabilities can improve genotype calls made from Illumina short reads in viruses with strong phylogenetic structure. We demonstrate that scaling the phylogenetic prior using a read-based maximum likelihood approach, equivalent to estimating the branch length of the connecting branch, improves results in both real and simulated data. The method yields a posterior probability for base calling combing read data information and phylogenetic information, that will allow users to balance the need for high base calling rates and high accuracy. We show that these improvements hold across sequencing conditions, and for different categories of sites that are typically problematic for either biological or technical reasons. We also note that in all analyses on real data, the gold standard treated as truth was the base calls obtained using standard methods based only on the read likelihood (state-of-the-art, or SA, pipeline) in high coverage data. Our analyses then evaluated performance in subsampled lower depth data. As such, our scheme for comparing the base-calling methods would, if anything, will be biased towards favoring the standard methods.

The improvement due to the use of a phylogenetic prior will, in general, depend on how representative the data used to infer the tree are for the sample data. Our analyses on real data included data that were published after the release of the database we used to generate the phylogenetic prior. One of the data sources, however, included older sequences that were likely well-represented by the database. Our study therefore includes both scenarios in which: (1) the database is incomplete and the new data includes strains, or substrains, not represented in the database, and (2) the database includes sequences closely related to the data being corrected. In the first scenario, if the viral sequencing reads originate from a highly divergent sequence, relative to the sequences in the database or the ancestral sequences implied by the tree, the scaling factor will estimate this mismatch and down-weight the evidence from the phylogenetic prior accordingly. As such, including the phylogenetic prior will not lead to reduced performance, even when the database does not contain representative sequences.

Despite its rapid evolution, SARS-CoV-2 provides a favorable setting for phylogenetically-informed genotype calling because of its extensive representation in public sequence databases. For less densely sampled viruses, the potential gains may be constrained by greater divergence between reference sequences and the genomes being analyzed, particularly at low sequencing depth. The broader applicability of this approach therefore depends on the availability of sufficiently representative reference sequences and the read evidence needed to accommodate departures from the phylogenetic prior.

## 6. Competing interests

No competing interest is declared.

## 7. Author contributions statement

M.A. and R.N. conceived the experiments and analyzed results. L.P. shared scripts and assisted with running analyses. All authors contributed to writing and reviewing the manuscript.

## 8. Acknowledgments

We gratefully acknowledge all data contributors, i.e. the Authors and their Originating laboratories responsible for obtaining the specimens, and their Submitting laboratories for generating the genetic sequence and metadata and sharing via the GISAID Initiative, on which this research is based. We also express our gratitude to the bioinformatics community who have curated and hosted data at pandemic scale, and to researchers who have shared their sequencing data through other public databases, especially the Sequencing Read Archive (SRA). We also thank the anonymous reviewers for their valuable suggestions.

## 9. Funding

This work used the Bridges-2 system at the Pittsburgh Supercomputing Center (PSC) through allocation BIO180028 from the Advanced Cyberinfrastructure Coordination Ecosystem: Services & Support (ACCESS) program. This work was supported by National Institutes of Health (NIH) grants [R00GM144747 to L.P., T32GM132022 to M.A.]. M.A. also received funding from the UC Berkeley Graduate Division’s Chancellor’s and Berkeley Connect fellowships.

## 10. Code Availability

Code to reproduce analyses and figures is available at https://github.com/marnieya/phylogenetic-base-calling.

## Notes

### Competing Interest Statement

The authors have declared no competing interest.

