## Supplemental Figures for "Error Correction of SARS-CoV-2 Genomic Sequences Using a Phylogenetic Prior"

**Supplementary Material for**: "Error correction of SARS-CoV-2 genomic sequences using a phylogenetic prior"

**S1. Accuracy and call rate variability in simulations**. Individual points represent simulated runs, each with a per-site mean error rate of 0.4%.

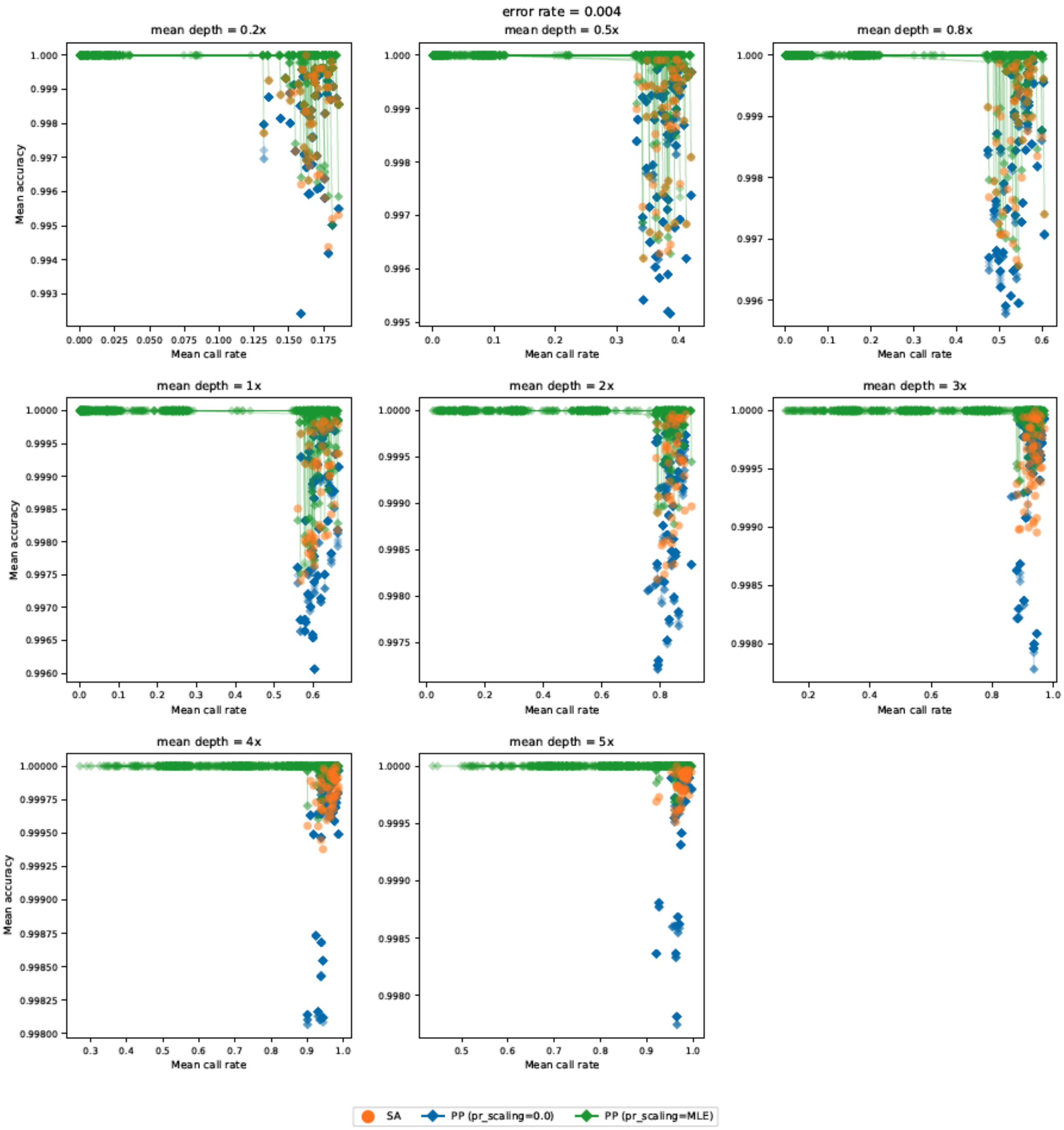

**S2. Accuracy and call rate variability in all real data.** Individual points represent each sequencing run, with metrics calculated over the entire genome.

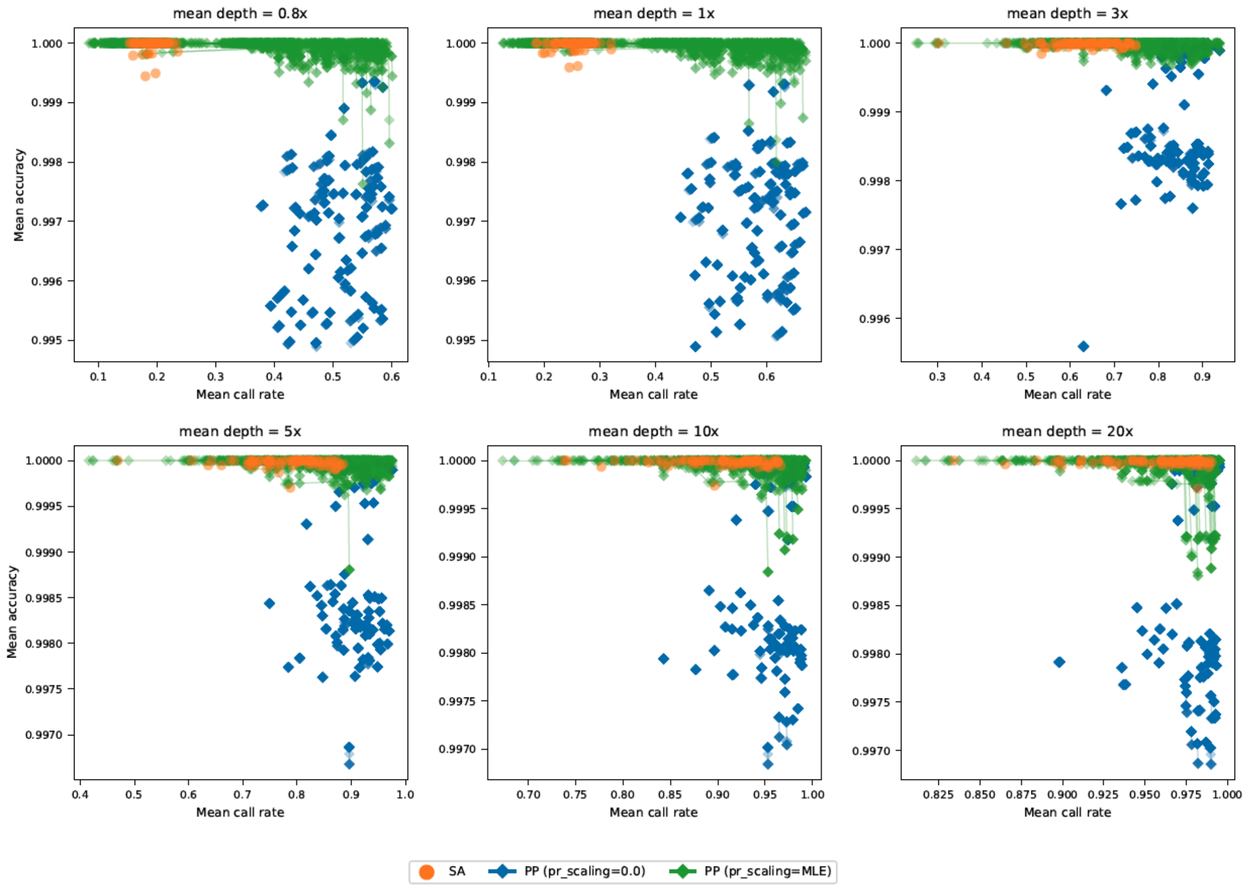

**S3. Per-position variability in depth and coverage.** Values are averaged across samples for all mean depths in simulated (top) and real (bottom) data.

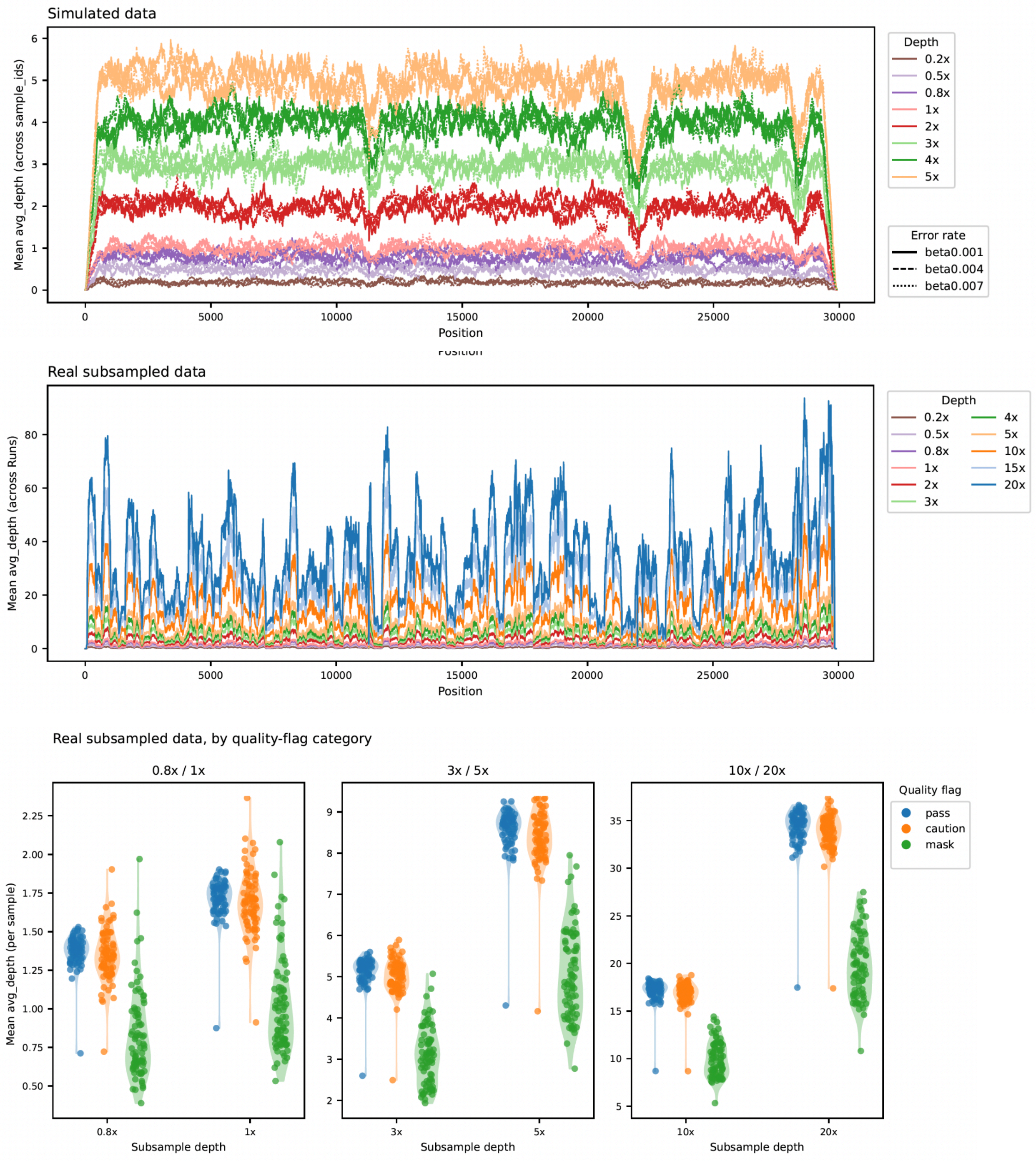

**S4. Call rate crossover thresholds.** We report the most stringent threshold possible where PP meets (but does not go below) SA mean call rate for all runs at each subsampled depth.

| Subsample mean depth | PP threshold | PP call rate | SA call rate | PP accuracy | SA accuracy |
|---|---|---|---|---|---|
| **0.8x** | 1-1e-11 | 0.1981 | 0.1922 | 0.999996 | 0.999973 |
| **1x** | 1-1e-11 | 0.2592 | 0.2527 | 0.999998 | 0.999973 |
| **3x** | 1-5e-12 | 0.6551 | 0.6345 | 0.999997 | 0.999984 |
| **5x** | 1-1e-12 | 0.7837 | 0.7833 | 0.999992 | 0.999976 |
| **10x** | 1-5e-12 | 0.908 | 0.902 | 0.999992 | 0.999982 |
| **20x** | 1-5e-12 | 0.9584 | 0.9563 | 0.999992 | 0.999982 |

**S5. Accuracy crossover thresholds.** We report the most relaxed threshold possible where PP meets (but does not go below) SA mean accuracy for all runs at each subsampled depth.

| Subsample mean depth | PP threshold | PP call rate | SA call rate | PP accuracy | SA accuracy |
|---|---|---|---|---|---|
| **0.8x** | 1-1e-5 | 0.5076 | 0.1922 | 0.999985 | 0.999973 |
| **1x** | 1-1e-5 | 0.5733 | 0.2527 | 0.999985 | 0.999973 |
| **3x** | 1-5e-5 | 0.8367 | 0.6345 | 0.999989 | 0.999984 |
| **5x** | 1-1e-4 | 0.9074 | 0.7833 | 0.99998 | 0.999976 |
| **10x** | 1-1e-5 | 0.9566 | 0.902 | 0.999983 | 0.999982 |
| **20x** | 1-1e-4 | 0.9795 | 0.9563 | 0.999983 | 0.999982 |
