## Supplementary material for "Error Correction of SARS-CoV-2 Genomic Sequences Using a Phylogenetic Prior": Dataset Supplement

### Supplementary Appendix

All genome sequences and associated metadata supporting the findings of this study can be accessed through the persistent digital object identifier https://doi.org/10.55876/gis8.260918ym

In addition to the minted DOI, GISAID also communicates the aggregation of GISAID accession numbers (EPI_ISL_IDs) through the corresponding EPI_SET_260918ym identifier to facilitate both, the acknowledgment of all data contributors and the direct retrieval of the underlying data from GISAID used in this study.

#### hCoV-19 Virus Data Summary

| GISAID Identifier | Digital Object Identifier | Number of individual viruses | Data Collection range | Number of countries/territories |
|---|---|---|---|---|
| EPI_SET_260918ym | https://doi.org/10.55876/gis8.260918ym | 69 | 2020-04-26 to 2023-08-28 | 13 |
